# Strain-dependent variation in maternal care and early behavioral development in laboratory rats

**DOI:** 10.1101/2025.11.05.686817

**Authors:** Grace E. Pardo, Alondra Casas Pary, María Esther Gutiérrez Ccori, Mabel Choque Aguilar, Lucero B. Cuevas, Enver M. Oruro, Luis F. Pacheco-Otalora

## Abstract

Understanding how maternal behavior varies among different laboratory rat strains is essential for improving the translational relevance of preclinical neurodevelopmental models. In this study, we compared maternal care patterns and the early development of offspring in three commonly used rat strains: Wistar, Sprague-Dawley (SD), and Spontaneously Hypertensive Heart Failure (SHHF). Maternal behaviors were recorded from postpartum day (PPD) 1 to 5 during both light and dark phases and analyzed using both conventional frequency-based methods and behavioral transition network analysis. Pup development was assessed from postnatal (PND) 6 to 22, including measures of somatic growth, eye-opening, reflex maturation, and ultrasonic vocalizations (USVs). We found significant strain differences in both the frequency and organization of maternal behaviors. SD dams exhibited reduced high-crouch nursing and fewer behavioral transitions across postpartum days. In contrast, SHHF dams spent more time in the nest without nursing and engaged in more frequent self-grooming, particularly during the dark phase. Network analysis revealed distinct transition patterns among strains, capturing qualitative differences in maternal dynamics not evident in conventional analysis. Strain differences also emerged in pup development. SHHF pups showed delayed eye opening, reduced body weight gain, and slower performance in several reflexes compared to Wistar and SD pups. Additionally, USV analyses revealed that SD and SHHF pups emitted fewer and shorter calls in both isolation-induced and maternal-potentiated contexts, especially in the low-frequency range. These findings underscore the importance of considering strain-specific profiles of maternal behavior and infant development when modeling early neurodevelopmental trajectories. SHHF rats may be particularly useful for studying early-life vulnerabilities relevant to human conditions associated with perinatal adversity. Moreover, behavioral transition networks offer a sensitive approach to reveal subtle differences in maternal caregiving strategies across strains.

## 1. INTRODUCTION

In altricial species such as rats, maternal behavior provides the primary source of sensory stimulation, physiological regulation, and protection during the early postnatal period. These maternal interactions are not only essential for immediate survival but also play a fundamental role in shaping the trajectory of neurobehavioral development in the offspring (Lucion & Bortolini, 2014; Mogi et al., 2011, 2017). Through behaviors such as nursing, licking, nest attendance, and contact maintenance, the dam regulates critical functions, including thermoregulation, emotional reactivity, and early learning (Esposito et al., 2015; Mota-Rojas et al., 2022). Several studies have demonstrated that maternal care is a dynamic and responsive process shaped by reciprocal interactions between mother and pup. For instance, pup vocalizations, posture, and movement can elicit specific maternal responses, creating a finely tuned system of sensory exchange (Febo et al., 2008; Nagasawa et al., 2012; Yoshida et al., 2013). These interactions help to establish the early mother-infant bond and contribute to the formation of the neural circuits involved in affective and social behavior later in life.

Research in rodent models has shown that naturally occurring variations in maternal care, particularly in licking/grooming and arched-back nursing, have long-lasting effects on offspring physiology and behavior. Offspring of highly licking/grooming mothers show reduced stress reactivity, improved cognitive performance, and increased hippocampal glucocorticoid receptor expression, effects that are mediated by epigenetic mechanisms (Champagne et al., 2003; D. Francis et al., 1999; Meaney, 2001). These behavioral styles are transmitted across generations not through genetics, but via the early postnatal environment, as demonstrated in cross-fostering studies (D. D. Francis et al., 2000, 2002). Moreover, maternal care impacts affective and social domains, including play behavior, social exploration, and anxiety-like responses, possibly through oxytocinergic signaling (Beery et al., 2016; Starr-Phillips & Beery, 2014). Despite this compelling evidence, most maternal care and early development studies have focused on a single rat strain, typically Long Evans or Sprague Dawley. This limited approach overlooks the potential contribution of strain-related genetic and physiological differences in shaping maternal care patterns. Several authors have emphasized the importance of considering naturally occurring variation in maternal care to improve the ecological validity and translational potential of rodent models (Beery & Francis, 2011). While such variation has been extensively studied within strains, less attention has been paid to inter-strain differences. However, recent work has highlighted that strain background and individual variability can significantly impact behavioral phenotypes and neurodevelopmental outcomes, underscoring their importance for reproducibility and interpretation in preclinical models (Löscher, 2024; Löscher et al., 2017). Strain differences may affect not only the expression of specific maternal behaviors but also their organization, responsiveness to pup cues, and capacity for adaptation under different physiological or environmental conditions.

A particularly underexplored aspect of strain variation involves models with preexisting physiological conditions such as hypertension. Spontaneously hypertensive rats, including the SHHF strain, are widely used in cardiovascular and metabolic research, yet little is known about their maternal behavior or its influence on offspring development (Youcef et al., 2014). Previous studies have shown that maternal hypertension can impair fetal growth and alter placental function (Johnston, 1995; Sharkey et al., 2001), while hormonal imbalances during pregnancy may influence the neuroendocrine regulation of maternal behavior postpartum (Sharkey et al., 2005; Small et al., 2016). From a translational perspective, these models are relevant for understanding how maternal health conditions may shape the early caregiving environment and, consequently, infant neurodevelopment (Swanson & David, 2015). In the case of SHHF rats, their combined hypertensive and metabolic vulnerability offers a particularly compelling model to explore the intersection of maternal physiology and caregiving behavior.

To address these gaps, we aimed to characterize the early postpartum maternal behavior and offspring developmental milestones in three rat strains commonly used in biomedical research: Wistar, Sprague Dawley (SD), and spontaneously hypertensive heart failure (SHHF). The study was conducted in a laboratory located at a high altitude (∼3300 meters above sea level), an environmental condition that can influence both maternal and offspring physiology. Since most published data are derived from sea-level conditions, the generation of reference profiles under high-altitude settings is essential for contextualizing behavioral findings and ensuring the appropriateness of cross-study comparisons.

Using an observational design, we analyzed maternal behavior during the first postpartum days and examined physical and behavioral developmental markers in male and female pups, including body weight, eye opening, sensory-motor reflexes, and ultrasonic vocalizations. In addition, we applied network analysis to evaluate the structure of behavioral transitions within maternal care, providing a dynamic view of how caregiving patterns are organized across strains. This study aims to establish a comparative baseline that can inform future research on strain selection, maternal-infant dynamics, and neurodevelopmental programming.

## 2. MATERIAL AND METHODS

### 2.1. Animals

Female and male Wistar, Sprague Dawley (SD), and spontaneously hypertensive heart failure (SHHF) rats (Charles River, USA), approximately 10-11 weeks old, were obtained from the animal facility of the Andean University of Cusco, located at 3300 meters above sea level. Animals were group-housed (4 per cage) in individually ventilated cages (30 x 30 x 18 cm), maintained under a 12h-light/dark cycle (lights on at 6:00 A.M.), with controlled temperature (22-23°C) and humidity (45%). Food and water were available *ad libitum*. All procedures were conducted in accordance with the NIH Guide for Animal Care and Use of Laboratory Animals and were approved by the Institutional Ethics Committee (CIEI) of the Andean University of Cusco (N°002-2022-CIEI-UAC).

### 2.2. Study procedure

After confirming four regular estrous cycles, 90-day-old virgin females (Wistar= 10 females; SD= 15 females; SHHF=11 females) were paired with males during the evening of proestrus. The presence of spermatozoids in the vaginal smear, verified under a light microscope the following morning, was taken as a positive indication of mating. This day was designated as gestational day 0 (GD) 0. Two SD females were excluded due to unsuccessful mating. Gestation was monitored daily, and body weight was recorded at GD 0, 5, 10, 15, 18, and 20, between 1:00 and 2:00 P.M. On GD 17-18, pregnant females were individually housed in transparent Plexiglas cages (30 x 30 x 18 cm) lined with clean shavings. These cages were monitored twice daily (9:00 A.M. and 5:00 P.M.) for signs of parturition. The day of birth was considered postpartum day (PPD) 0. Three SHHF females were excluded at this stage due to miscarriages. On the morning of PPD 1, litter sizes were standardized to 7 pups for Wistar, 10 for SD rats, and 8 for SHHF dams, ensuring an equal proportion of male and female pups whenever possible. Five females were excluded at this point (2 Wistar, 2 SD, 1 SHHF) due to insufficient number of pups. During the standardization, the pups were weighed, and the dam and litter were transferred to a clean cage with fresh bedding. Cages were not changed again until PPD 6.

Maternal behavior was recorded daily from PPD 1 to PPD 5. Assessments of the pups’ somatic development and reflexes began on PPD 6, between 3:00 P.M. and 5:00 P.M. Prior to testing, dams were gently removed from their home cages and placed in clean, bedding-lined holding cages within the same room. The home cage containing the pups was then transported to a temperature- (27–30°C) and humidity-controlled (40%) testing room. Each pup was marked daily on the back with a non-toxic indelible marker before testing. All assessments were video recorded and performed in the following sequence: weighing, evaluation of eye-opening, paw grasping, superficial righting, negative geotaxis, accelerated righting, startle response, and ultrasonic vocalizations. All pups in each litter were tested individually. After testing, the pups were returned to the animal facility room, and the dam was returned to her original home cage and cage rack. For all strains, the dam and pups remained in the same cage until weaning on postnatal day (PND) 23. During this period, cage changes were performed every four days.

### 2.3. Maternal behavior observation

Maternal behavior was assessed following the methodology described in our previous studies (G. E. Pardo et al., 2024; G. V. E. Pardo et al., 2016). Each dam was observed in her home cage for a 72-minute period, comprising two sessions during the light phase (9:00 A.M. and 2:00 P.M.) and one session during the dark phase (6:00 P.M.) from postpartum day (PPD) 1 to PPD 5. Observations during the dark phase were conducted under red light to minimize disturbance. During each 72-minute session, maternal behavior was scored every 3 minutes, resulting in 25 observations per session and 75 observations per day per dam. The following maternal behaviors were recorded: high crouch posture (HG, mother nursing pups in an arched-back posture); low crouch posture (LW, mother nursing pups in a “blanket” low arched back posture); supine posture (SUP, a passive posture in which the mother is lying on her back or side while the pups nurse); licking the pups (L, licking/grooming the surface of their bodies and their anogenital regions); HG posture at the same time the mother licking the surface of pup’ bodies and their anogenital regions (HG/L); dam in the nest (DN; dam in the nest area, mostly besides the pups, without showing any nursing behavior), dam licking the pups (L), dam recovering pups (R), nest building (NB), mother off the nest (OFF, the lactating female is out of the nest), self-grooming (SG), and pup off the nest (PO). On PPD 1, the 9:00 A.M. observation was conducted at least 30 minutes after transferring the dam and her litter to new cages to allow for acclimatization to the environment. The analysis focused on the number of events recorded for each behavioral category. Additionally, we quantified the total number of maternal behavioral transitions (i.e., shifts from one behavior to another) for each dam during the light and dark phases. Data are presented as percentages of behavioral events, averaged across the two light-phase sessions and separately for the dark-phase session.

### 2.4. Physical and reflex development assessment

Developmental milestones and reflexes were assessed in all pups from the same litter used for maternal observations. To avoid disturbing the nest and interfering with maternal care, testing began at PND 6, one day after the maternal observation period ended. Each litter was treated as an experimental unit. Performance scores were calculated as the average of values obtained from all pups of the same sex within a litter, allowing for the separate analysis of male and female data.

#### 2.4.1. Body weight gain

Each pup was weighed individually on PND 1, and body weight was monitored daily from PND 6 to PND 22 during the developmental assessment period.

#### 2.4.2. Eye-opening

Eye-opening was assessed daily from PND 10 to PND 17. Each pup was gently held while the eyelids were visually inspected. The scoring system, based on previous studies (Demaestri et al., 2020; Pardo et al., 2023), was as follows: both eyes closed= 0, half eye open =0.5, and both eyes half open =1; one eye fully open and another eye half open =1.5; and two eyes fully open =2.

### 2.5. Reflex developmental assessment

A standard battery of reflex assessments was used, based on previous protocols (Baharnoori et al., 2012; Heyser, 2003; Nguyen et al., 2017). The initiation and termination of each reflex test varied according to the expected appearance and disappearance windows of the reflex in early development.

#### 2.5.1. Paw grasping

Assessed from PND 6 to PND 10. Pups were gently lifted by the torso to allow the limbs to hang freely. A blunt metal rod was gently rubbed across the palm of each forelimb and hindlimb. A successful reflex was defined by grasping the rod on two consecutive days. Scoring was: no grasping=0, successful grasping by one forepaw or hindpaw=1, successful grasping by both forepaws or hindpaws=2. For each pup’s paw grasping reflex, the scores for the forelimb and hindlimb reflexes were averaged.

#### 2.5.2. Superficial righting

Tested from PND 6 to PND 9. Pups were placed on their backs on a flat surface, and the time taken to right themselves (return to four paws) was recorded.

#### 2.5.3. Negative geotactic reaction

Assessed from PND 6 to PND 9. Pups were placed head-downward on a 30° inclined surface. The latency to rotate and orient their heads upward was recorded as a measure of geotactic response.

#### 2.5.4. Accelerated righting

Tested from PND 13 to PND 16. Pups were gently dropped from a height of 40 cm in a supine position above a foam pad. Successful righting was scored as follows: pup landing on its back=0, pup falling on its side (left or right) =1, and pup landing on its paws=2.

#### 2.5.5. Auditory startle

Conducted from PND 10 to PND 14. A small bell was rung 30 cm above the pup. Observable startle responses such as body jerks, kicking, squirming, or a combination of those behaviors were recorded. Responses were scored as follows: 1 = presence of startle, 0= no response.

### 2.6. Maternal potentiation of ultrasonic vocalizations (USVs)

On PND 6, 9, and 14, all pups from each litter were tested for isolation-induced and maternal-potentiated ultrasonic vocalizations (USVs) following a previously established protocol (Hofer et al., 2001). USV testing was conducted 15 minutes after the completion of the reflex assessments. Each pup was transported individually to a separate testing room maintained at 24–25 °C. The pup was placed inside a 500 mL plastic beaker, and USVs were recorded during a 2-minute isolation period. A condenser microphone (UltraSoundGate CM16; Avisoft Bioacoustics, Berlin, Germany) was positioned 12 cm above the center of the beaker. The microphone was connected to a computer via an UltraSoundGate 116 USB audio device and was controlled using Avisoft Recorder 2.7 software. The system had a sampling rate of 250 kHz and a frequency detection range of 20–100 kHz. Immediately after the first isolation period (ISO 1), the pup was placed in a holding cage containing its awake dam for 1 minute, allowing direct contact. The pup was then returned to the testing room for a second 2-minute isolation session (ISO 2), during which USVs were recorded again under the same conditions. After testing, each pup was returned to its home cage, and the procedure was repeated for the next pup. Between tests, the testing chamber and transport container were cleaned with distilled water to avoid odor contamination.

Spectrogram analysis of the USV recordings was performed using Avisoft-SASLab Pro software. Each recording file was manually reviewed to ensure the accuracy of automated call detection. A call was defined as a continuous, uninterrupted vocalization. For each identified call, the following parameters were analyzed: peak frequency (kHz), onset time (s), and offset time (s). Additionally, we computed: peak frequency (kHz), start time (s), and end time (s). We calculated the number of low-frequency calls (number of calls < 60 kHz) and the number of high-frequency calls (number of calls > 60 kHz), duration of calls of low and high-frequency calls. Call frequency distributions were visualized using probability density functions (PDFs). For statistical analyses, the mean values of each USV parameter were calculated per pup and averaged across pups of the same litter.

### 2.7. Statistical analysis

Statistical analyses were conducted using GraphPad Prism 9.1.2 software (GraphPad, San Diego, CS, USA). Maternal behavior and developmental behavioral data were analyzed using two-way repeated measures analysis of variance (RM ANOVA), followed by Šídák’s multiple comparisons post hoc test. For the analysis of ultrasonic vocalization (USV) characteristics, differences in peak frequency and other continuous variables were evaluated using either unpaired t-tests (for normally distributed data) or Mann–Whitney U tests (for non-parametric comparisons). Results are expressed as mean ± standard error of the mean (SEM) or as median with interquartile range (25th– 75th percentile), depending on data distribution. Statistical significance was set at *p* < 0.05. In figures and tables, asterisks indicate significant group differences.

### 2.8. Network analysis

As a complement to the statistical analysis, we applied network analysis to visualize and characterize qualitative changes in maternal behavioral transitions across the observation period. The methodology follows the procedures previously described in our earlier work (Pardo et al., 2024). For each dam, behavioral transition data were averaged separately for the light phase (based on observations at 9:00 A.M. and 2:00 P.M.) and the dark phase (6:00 P.M.). Individual networks were constructed per dam and subsequently averaged within each strain group. In the network representation, nodes correspond to six defined maternal behavior components: HG, HG/L, LW, SUP, DN, and OFF. Links represent transitions between behavioral states. Arrows indicate the direction of each transition, and the thickness of each link reflects the frequency of that specific transition within the observation window.

## 3. Results

### 3.1. Maternal behavior

Maternal behavior was compared among Wistar, SD, and SHHF rat strains from PPD 1 to PPD 5, using two-way repeated measures ANOVA for both light and dark phases. No significant differences were found between observations taken at 9:00 A.M. and 2:00 P.M.; therefore, these were averaged and analyzed together to represent the light phase. Comparative analyses revealed strain-specific differences in maternal care profiles (see **Figure 1**), and a summary of the statistical results is presented in **Table 1**.

**Figure 1.**
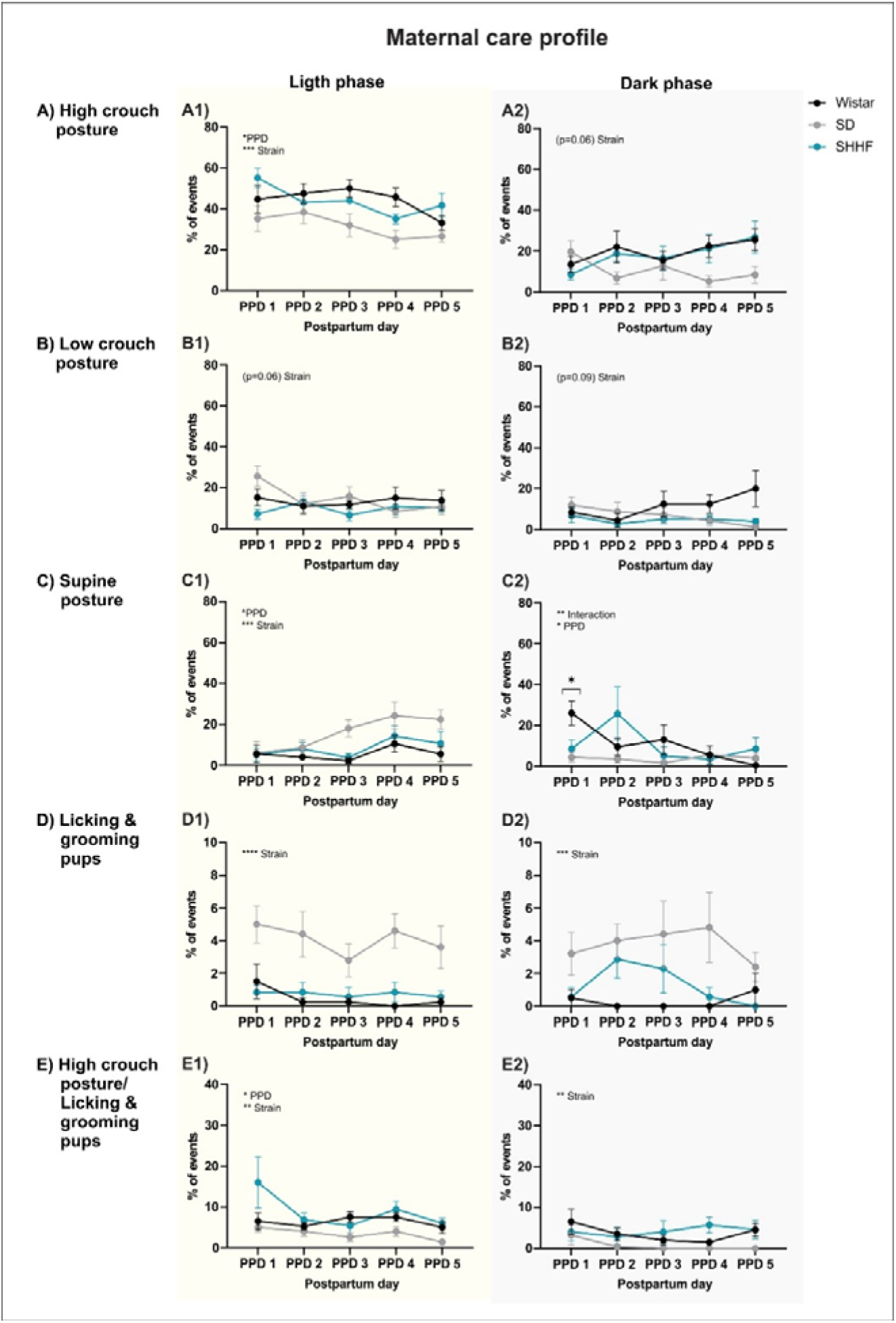
Maternal behavioral profile of Wistar, SD, and SHHF rat strains during early postpartum days (PPD 1-5). **A**) High crouch nursing posture (HG) was significantly lower in SD dams compared to the other strains during both the light (**A1**) and dark phases (**A2**). **B**) Low crouch nursing posture (LW) showed comparable frequencies across strains during both phases, with minor differences (**B1,B2**). **C**) Supine posture (SUP) increased progressively in SD dams during the light phase (**C1).** During the dark phase (**C2**), post hoc analysis revealed fewer SUP events in SD and SHHF dams compared to Wistar dams on PPD 1.**D**) Licking and grooming of pups (L) was more frequent in SD dams across postpartum days during both the light (**D1**) and dark phases (**D2**), particularly in comparison to Wistar and SHHF dams. **E**) The combination of high crouch posture with simultaneous licking and grooming (HG/L) was reduced in SD dams relative to the other strains across both phases (**E1, E2**). Data represent mean ± SEM (n = 7-10 per strain). Two-way ANOVA was used for statistical analysis. Asterisks indicate significant effects; see **Table 1** for details. \**p*<0.05, \*\**p*<0.01, \*\*\**p*<0.001, \*\*\*\**p*<0.0001.

High crouch (HG) posture was significantly reduced in SD dams compared to Wistar and SHHF dams during both the light (**Figure 1, A1**) and dark phases (**Figure 1, A2**). Low crouch (LW) posture showed only modest strain variation. During the dark phase, LW was more frequent in Wistar dams than in the other two strains (**Figure 1, B2**), while no clear differences were observed during the light phase (**Figure 1, B1**). Supine posture (SUP) was more prevalent in SD dams during the light phase (**Figure 1, C1**). In contrast, during the dark phase, this behavior increased significantly only in Wistar dams on PPD 1 (**Figure 1, C2**). Licking and grooming (L) behavior also differed among strains. SD dams exhibited significantly higher levels of pup-directed licking than the other strains during the light phase (**Figure 1, D1**), and this behavior remained higher in SD and SHHF dams during the dark phase, although SHHF levels were still lower than those of SD (**Figure 1, D2**). The HG/L behavior, combining nursing in high crouch posture with simultaneous licking, was reduced in SD dams compared to Wistar and SHHF across both phases (**Figure 1, E1 and E2**).

Additional maternal behaviors are shown in **Supplementary Figure S1**. Dam off-nest (OFF) behavior did not differ significantly between strains during the light phase (**Figure S1, A1**). However, during the dark phase, SHHF dams showed increased nest-leaving on PPD 1, and SD dams exhibited a similar increase on PPD 5 (**Figure S1, A2**). Dam in-nest (DN) behavior showed significant variation across strains. SD dams spent more time in the nest without active nursing than Wistar and SHHF dams during the light phase (**Figure S1, B1**), and this was particularly pronounced on PPD 1 during the dark phase (**Figure S1, B2**). Nest-building (NB) behavior did not differ significantly among strains during the light phase (**Figure S1, C1**), but SD dams showed a slight increase during the dark phase (**Figure S1, C2**). Self-grooming (SG) was more frequent in SHHF dams across both phases, especially on PPD 1, 4, and 5 during the dark phase (**Figure S1, D1 and D2**). Finally, behavioral transitions, defined as the total number of switches between maternal behavior components, differed by strain: SD dams exhibited fewer transitions than Wistar and SHHF dams in both the light and dark phases (**Figure S1, E1 and E2**).

### 3.2. Maternal behavior transition profile

To explore qualitative differences in maternal behavioral organization among the three rat strains, we conducted a network-based transition analysis during the light and dark phases of the first postpartum days (**Figures 2 and 3**), complemented by heatmaps illustrating the strength of specific behavioral sequences (**Figure S2**). These tools provide a dynamic caregiving perspective by visualizing how frequently dams shift between different maternal behaviors, beyond absolute frequencies.

**Figure 2.**
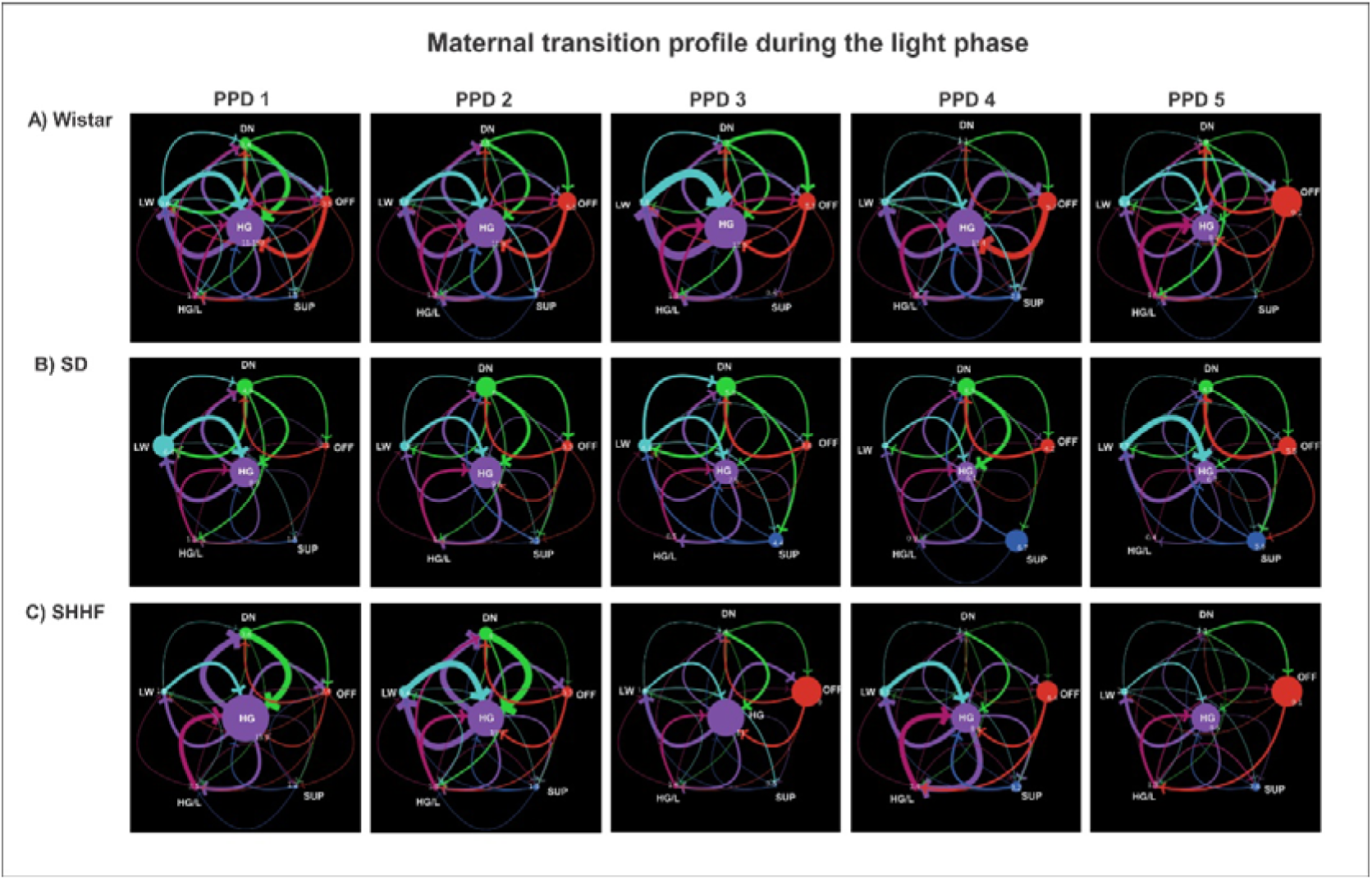
Maternal behavioral transition networks during the light phase across postpartum days (PPD 1–5) in three rat strains. Transition networks illustrate changes in maternal behavior dynamics in Wistar (**A**), Sprague-Dawley (SD) (**B**), and Spontaneously Hypertensive Heart Failure (SHHF) (**C**) dams. Each network is composed of six nodes representing distinct maternal behaviors: HG (high crouch posture), HG/L (high crouch with simultaneous licking and grooming), LW (low crouch posture), SUP (supine posture), DN (dam in the nest without nursing), and OFF (dam off the nest). Node size reflects the average frequency of each behavior, while arrows indicate transitions between behaviors. Thicker arrows represent more frequent transitions. Networks were constructed by averaging data from light-phase observations (9:00 A.M. and 2:00 P.M.) across PPD 1 to 5 (n = 7-10 dams per strain). Graphs were generated using NetLogo software.

**Figure 3.**
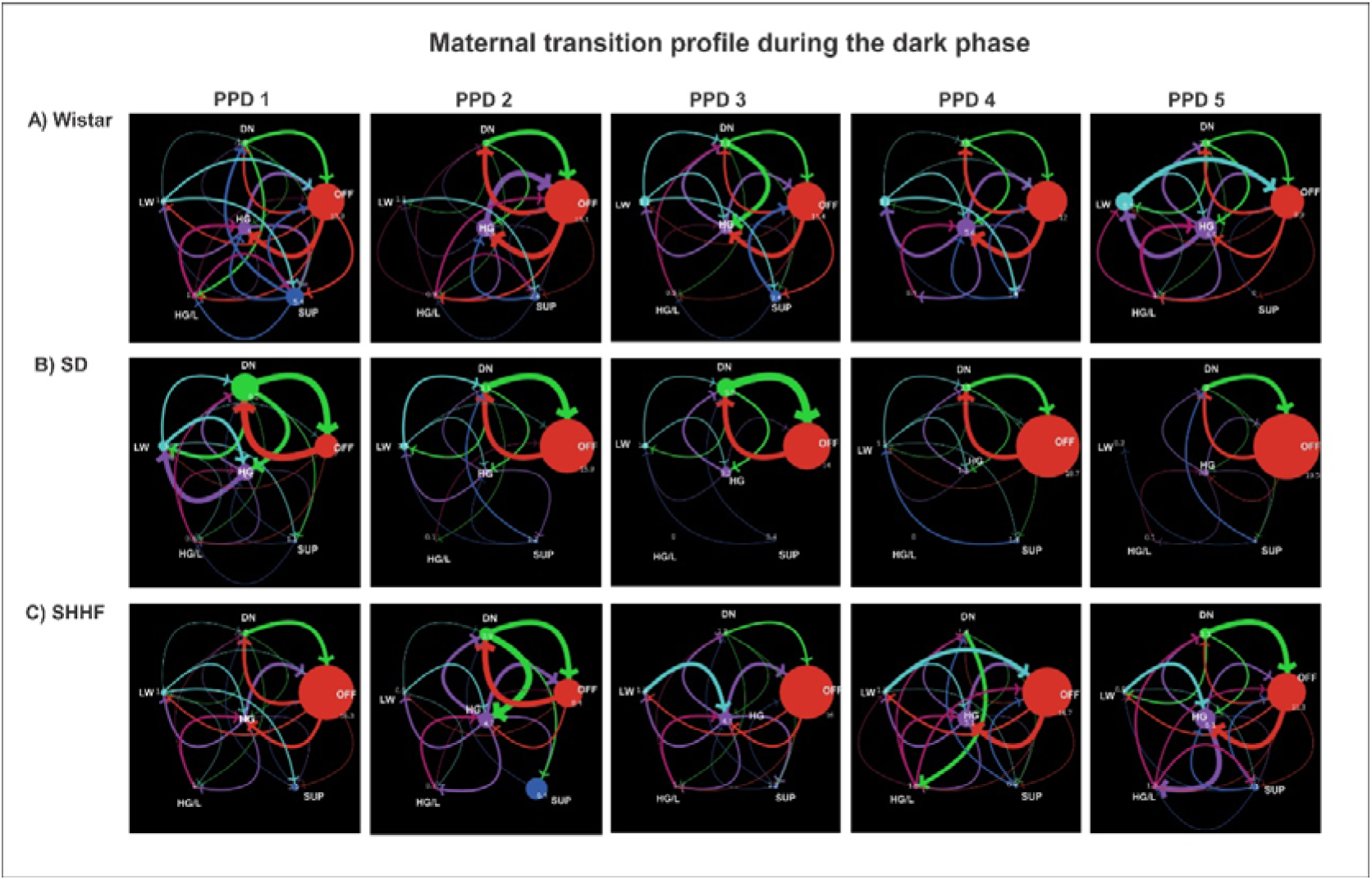
Maternal behavioral transition networks during the dark phase across postpartum days (PPD 1–5) in three rat strains. Transition networks illustrate changes in maternal behavior dynamics in Wistar (**A**), Sprague-Dawley (SD) (**B**), and Spontaneously Hypertensive Heart Failure (SHHF) (**C**) dams. Each network includes six nodes representing distinct maternal behaviors: HG (high crouch posture), HG/L (high crouch posture with simultaneous licking/grooming), LW (low crouch posture), SUP (supine posture), DN (dam in the nest without nursing), and OFF (dam off the nest). Node size reflects the average frequency of each behavior, and directed arrows indicate transitions between behaviors. Thicker arrows denote more frequent transitions. Networks were generated by averaging data from dark-phase observations (6:00 P.M.) across PPD 1 to 5 (n = 7-10 dams per strain). Graphs were created using NetLogo software.

During the light phase (**Figure 2A**), Wistar and SHHF dams exhibited networks characterized by prominent HG (high crouch nursing posture) nodes, with frequent transitions to and from DN (dam in nest) and OFF (out of nest), indicating dynamic alternation between active caregiving and brief exits from the nest. In contrast, SD dams displayed more centralized networks with dense transitions between HG and HG/L (licking during high crouch) and fewer links involving OFF behaviors, suggesting a tendency to remain in the nest and sustain caregiving activity.

As the days progressed, SHHF dams maintained robust transitions between HG and HG/L, while Wistar networks increasingly incorporated OFF transitions. SD dams maintained a relatively stable transition profile with less pronounced OFF behavior across days. During the dark phase (**Figure 3**), overall behavioral activity increased in all strains, with OFF nodes becoming more prominent, especially in Wistar and SD dams, indicating more frequent nest exits. Wistar dams exhibited structured transition sequences such as HG → DN → OFF, while SHHF dams showed a more distributed network, with transitions spread across multiple behavioral states. SD dams, on the other hand, maintained a similar structure observed during the light phase but a slight increase in OFF-related transitions. Heatmaps (**Figure S2**) further highlight strain differences in the temporal organization of behavior sequences. Wistar dams consistently showed HG→DN and DN→OFF transitions in both phases. SD dams exhibited reduced variability, with fewer transitions involving OFF or DN states, whereas SHHF dams displayed variable transitions distributed across multiple states, including frequent transitions from DN and HG to OFF, particularly as postpartum days progressed. This pattern reflects a more dispersed behavioral structure rather than a consistently diverse nursing repertoire.

Together, these network patterns suggest strain-specific caregiving styles, where Wistar dams showed a distributed and alternating caregiving style with frequent behavioral shifts, SD dams maintained a focused, sustained caregiving profile with limited nest exits, and SHHF dams adopted a mixed strategy with marked variability and shifting sequences across postpartum days.

### 3.3. Pup’s developmental milestones

#### 3.3.1. Body weight gain and eye-opening

Strain differences in early physical development are shown in **Figure 4**. SHHF pups of both sexes exhibited reduced body weight gain compared to Wistar and SD pups (**Figure 4A**). A two-way repeated measures ANOVA with strain and PND as factors revealed a significant interaction (see **Table 2)**. Post-hoc analysis showed that SHHF males had significantly lower body weight from PND 10 onward (**Figure 4A1**), while SHHF females differed from the other strains as early as PND 6 (**Figure 4A2**). These growth differences persisted until PND 22 for both sexes.

**Figure 4.**
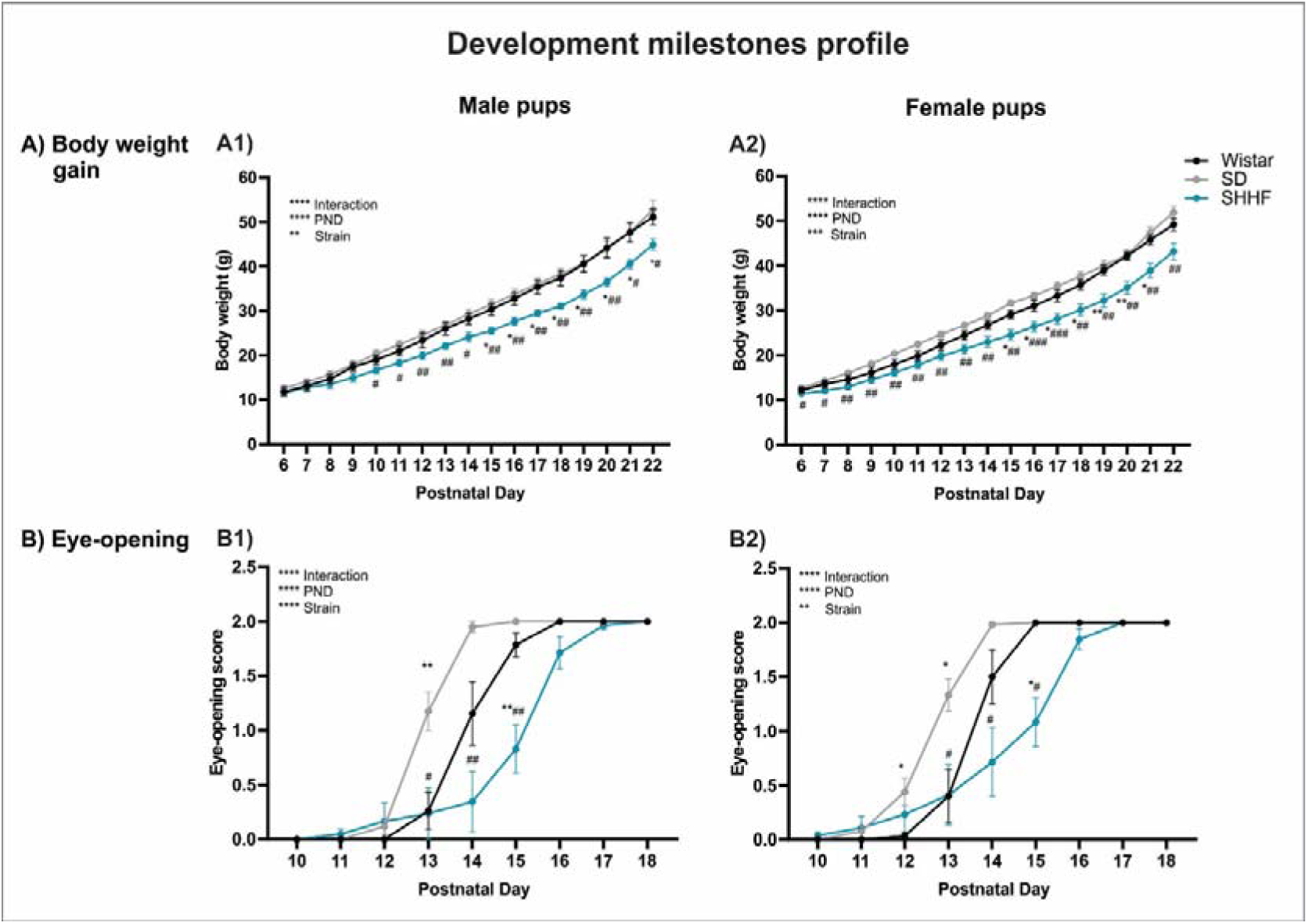
Developmental profiles of body weight gain and eye-opening in male and female pups from three rat strains. **A**) Growth trajectories for male (**A1**) and female (**A2**) pups show that SHHF pups consistently displayed reduced body weight across postnatal days (PND 6–22) compared to Wistar and SD pups. (**B**) Eye-opening scores for male (**B1**) and female (**B2**) pups indicate delayed eye-opening in SHHF pups relative to both Wistar and SD strains. Additionally, Wistar pups exhibited later eye-opening compared to SD pups. Data are presented as mean ± SEM (n = 7–10 per strain) and were analyzed using two-way ANOVA. Statistical details are reported in **Table 2**. \**p* < 0.05, \*\**p* < 0.01, \*\*\**p* < 0.001, \*\*\*\**p* <0.0001.

Eye-opening scores also differed significantly by strain and age, with a significant interaction between factors (**Table 2**). Post hoc analyses revealed that SHHF males delayed eye-opening compared to SD pups on PND 13, 14, and 15, and to Wistar pups on PND 15. Wistar males also scored lower than SD pups on PND 13 (**Figure 4B1**). In females, SHHF pups scored lower than SD pups on PND 13-15 and than Wistar pups on PND 15. Wistar females also had lower scores than SD pups on PND 12 and 13 (**Figure 4B2**).

#### 3.3.2. Reflex development

Reflex development was evaluated in male and female pups across postnatal days using two-way repeated measures ANOVA. Results are shown in **Figure 5**, with statistical details summarized in **Table 2**.

**Figure 5.**
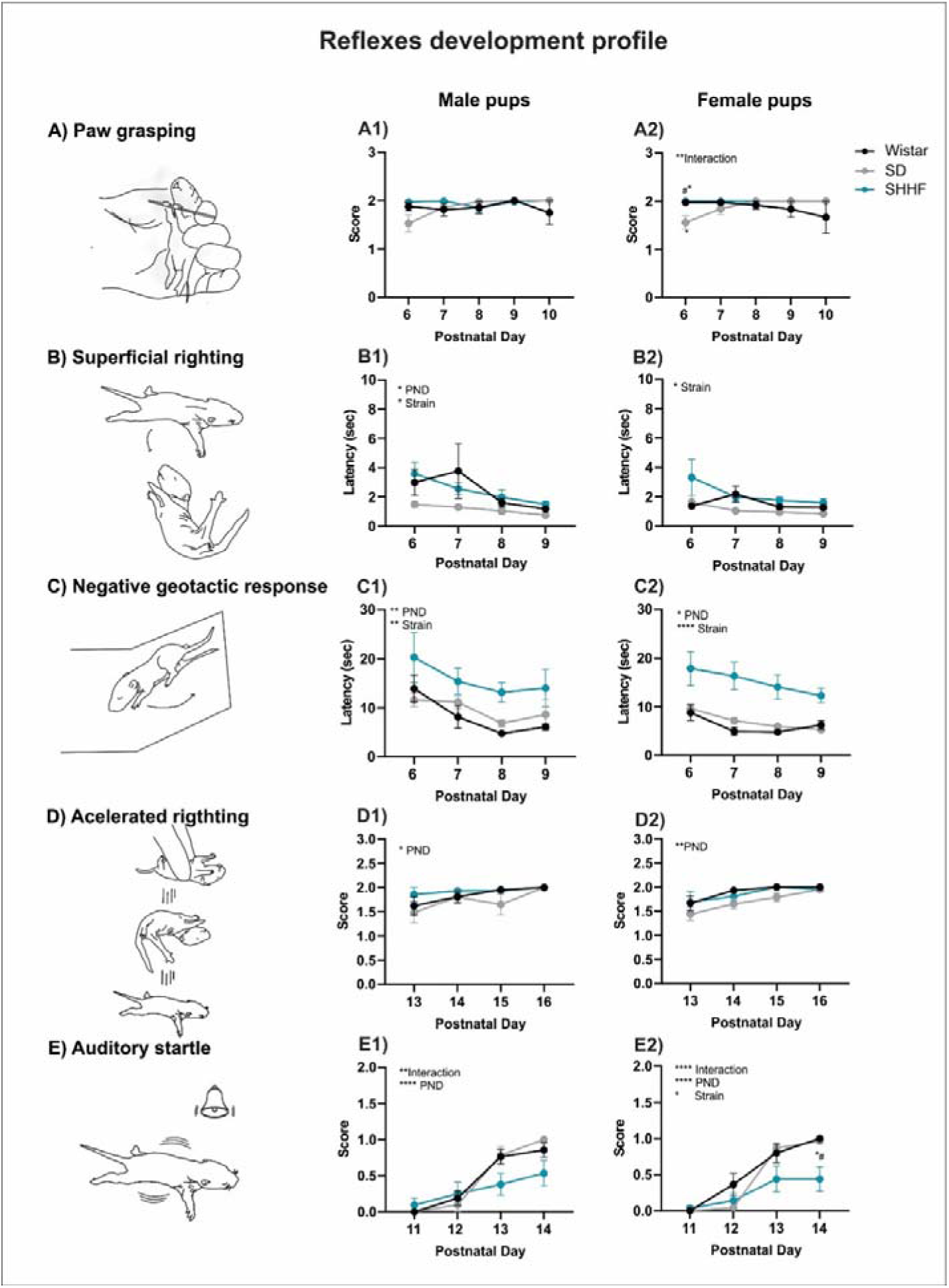
Reflex development profiles of male and female pups from Wistar, SD, and SHHF strains. **A**) Paw grasping reflex scores in male pups (**A1**) showed no significant differences among strains. In female pups (**A2**), SD rats exhibited lower scores on PND 6 compared to Wistar and SHHF pups. **B**) Superficial righting latency decreased over time in all groups. SD pups of both sexes (**B1** and **B2**) showed shorter latencies to righting compared to the other strains. **C**) Negative geotactic response latency was consistently longer in SHHF pups of both sexes (**C1** and **C2**), indicating delayed development of this reflex relative to Wistar and SD pups. **D**) Accelerated righting scores increased with age in all strains and sexes, with no strain-specific differences (**D1** and **D2**). **E**) Auditory startle responses showed a strain × age interaction. SHHF pups (**E1** and **E2**) exhibited reduced startle responses on PND 13–14 compared to Wistar and SD pups. Data are presented as mean ± SEM (n = 7–10 per strain), analyzed using two-way ANOVA. Detailed statistics are provided in **Table 2**. \**p* < 0.05, \*\**p* < 0.01, \*\*\**p* < 0.001, \*\*\*\**p* <0.0001.

Grasping reflex was assessed from PND 6 to PND 10 (**Figure 5A**). In males, no significant effects of strain, day, or their interaction were found (**Figure 5A1**). In females, a significant strain x PND interaction was found. Post hoc comparison revealed that SHHF and Wistar females performed better than SD females on PND 6 (**Figure 5A2**).

Superficial righting reflex was assessed from PND 6 to PND 9 (**Figure 5B**). In males, both strain and postnatal day had significant effects (**Figure 5B1**). In females, a significant main effect of strain was found (**Figure 5B2**). In both sexes, SD pups exhibited shorter righting latencies than Wistar and SHHF pups.

Negative geotactic response was evaluated from PND 6 to PND 9 (**Figure 5C**). In both males (**Figure 5C1**) and females (**Figure 5C2**), significant effects of strain and postnatal day were found. SHHF pups showed longer latencies to reorient uphill, indicating a delayed response compared to the other strains.

Accelerated righting was assessed from PND 13 to PND 16 (**Figure 5D**). In both sexes (**Figures 5D1 and 5D2**), performance improved significantly across postnatal days, but no strain differences were found.

Auditory startle response was assessed from PND 11 to PND 14 (**Figure 5E**). In males (**Figure 5E1**) and females (**Figure 5E2**), significant effects of strain, postnatal day, and their interaction were found. SHHF pups consistently exhibited lower startle scores, with significantly attenuated responses on PND 14 compared to Wistar and SD pups.

#### 3.3.3. USV calls

A two-way ANOVA was used to analyze ultrasonic vocalization (USV) patterns in male and female pups from the three strains at PND 6, 9, and 14. Results for isolation-induced USVs (ISO 1) are presented in **Figure 6**, and those for maternal potentiation USVs (ISO 2) are shown in **Figure 7**. Detailed statistical results are summarized in **Table 3**.

**Figure 6.**
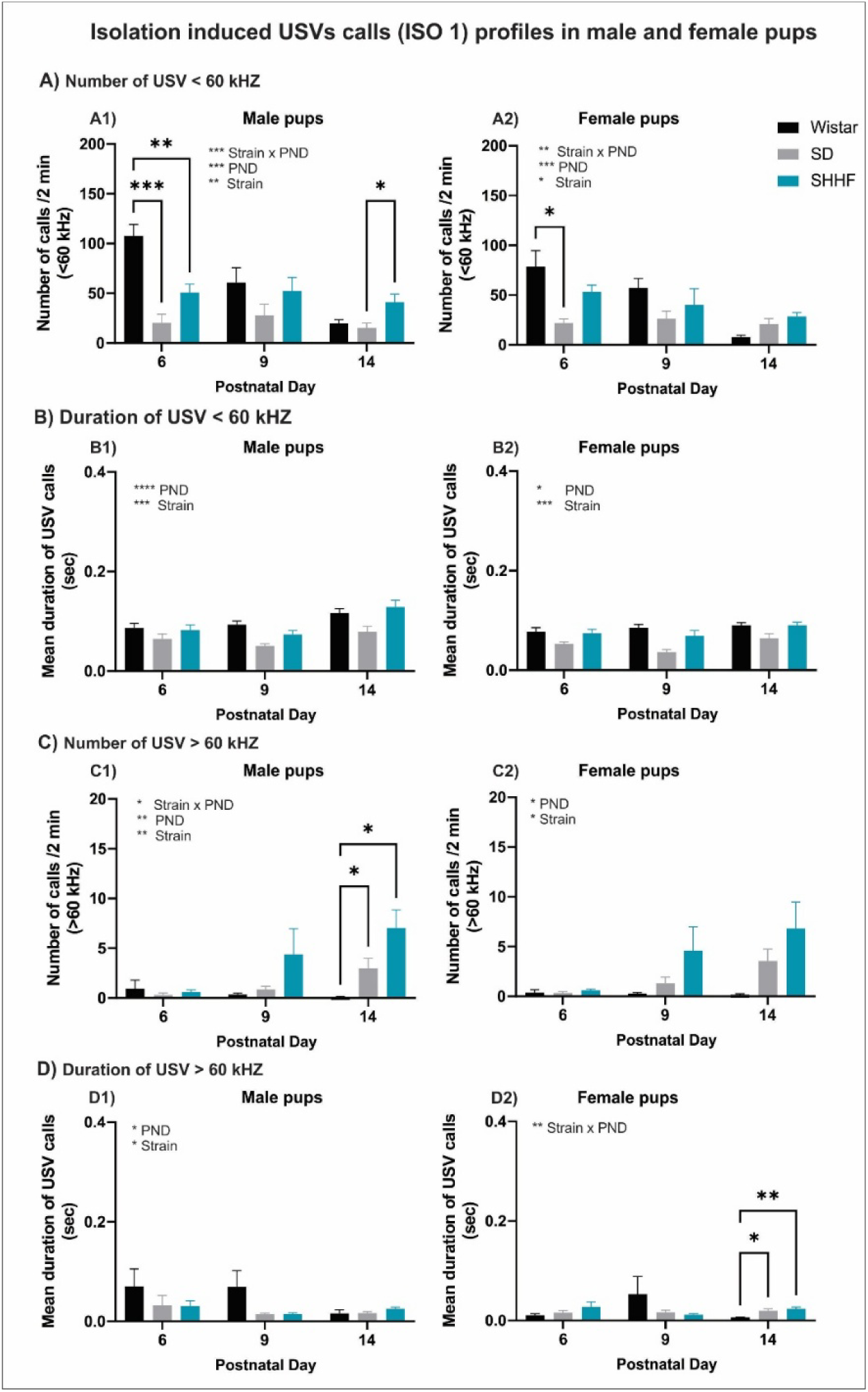
Profiles of isolation-induced ultrasonic vocalizations (USV) calls (ISO 1) in in male and female pups from three rat strains. **A**) Number of low-frequency USVs (< 60 kHz) in male (**A1**) and female (**A2**) pups was significantly reduced in SD and SHHF strains compared to Wistar pups, particularly on PND 6. On PND 14, SHHF male pups emitted more calls than SD pups. **B**) Duration of low-frequency USVs in males (**B1**) and females (**B2**) showed a main effect of strain and postnatal day, with SD pups emitting shorter-duration calls across ages compared to the other strains. **C**) Number of high-frequency USVs (> 60 kHz) increased with age in all groups. SHHF male pups (**C1**) emitted significantly more high-frequency calls at PND 14 than Wistar and SD pups. In females (**C2**), a strain and postnatal day effect was also observed, with SHHF pups tending to emit more calls. **D**) Duration of high-frequency USVs showed a main effect of age and strain in males (**D1**), with a general reduction in call duration across development. In females (**D2**), a significant strain × age interaction was found; post hoc tests revealed that SHHF and SD pups emitted longer calls than Wistar pups on PND 14. Data are presented as mean ± SEM (n = 7–10 per strain), analyzed using two-way ANOVA. Full statistics are available in **Table 3**. \**p* < 0.05, \*\**p* < 0.01, \*\*\**p* < 0.001, \*\*\*\**p* <0.0001.

**Figure 7.**
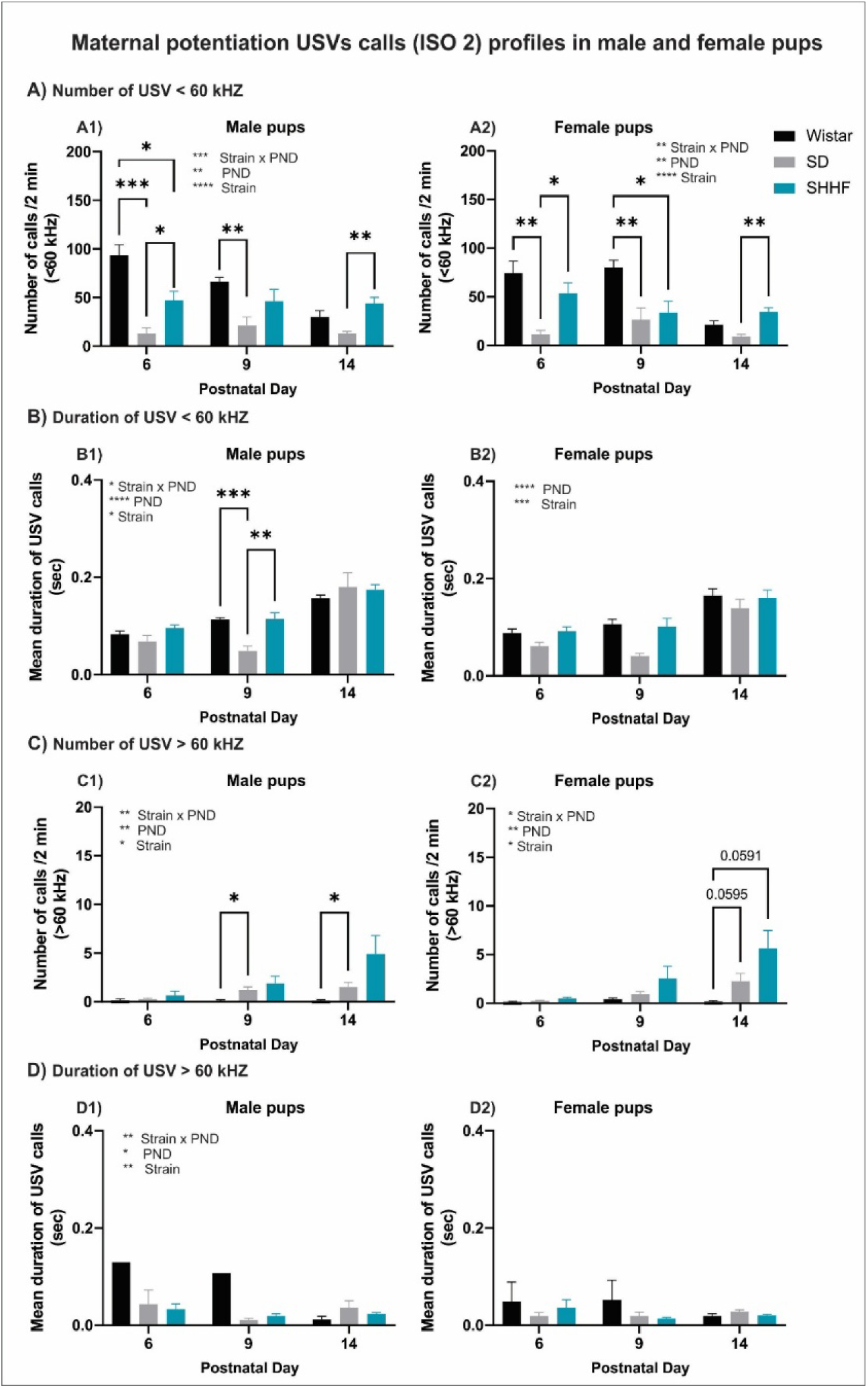
Profile of maternal potentiation ultrasonic vocalization (USV) calls (ISO 2) in male and female pups from three rat strains. **A**) Number of low-frequency USVs (<60 kHz) in males (**A1**) and females (**A2**) across postnatal days (PND) 6, 9, and 14. Wistar pups emitted significantly more USVs than SD and SHHF pups, with clear strain and developmental effects. **B**) Duration of low-frequency USVs (<60 kHz) in males (**B1**) and females (**B2**). Male SHHF pups showed significantly longer USV durations at PND 9 compared to the other strains. Female pups displayed a gradual increase in USV duration over time, with no major strain differences. **C**) Number of high-frequency USVs (>60 kHz) in males (**C1**) and females (**C2**). Male SHHF and SD pups produced more calls than Wistar pups, especially on PND 9 and 14. A similar pattern was observed in females, with marginal strain differences emerging on PND 14. **D**) Duration of high-frequency USVs (>60 kHz) in males (**D1**) and females (**D2**). Male SHHF and SD pups exhibited shorter durations than Wistar pups, particularly on early PNDs. No consistent differences were observed among female pups. Data are presented as mean ± SEM (n = 7–10 per strain). Two-way ANOVA revealed significant main effects of strain, postnatal day (PND), and their interaction. Statistical details are provided in **Table 3**. \**p* < 0.05, \*\**p* < 0.01, \*\*\**p* < 0.001, \*\*\*\**p* <0.0001.

During the first isolation (ISO 1), pups from all strains emitted USV primarily in the 40–45 kHz range (<60 kHz), which is typically associated with distress signaling and thermoregulatory needs. In addition, SD and SHHF pups also produced calls with higher peak frequencies (75–80 kHz) (>60 kHz), suggesting a broader spectral range of vocalization at these ages. The distribution of frequency peaks during both isolation contexts is illustrated in **Supplementary Figure S3**.

##### Isolated-induced USV calls (ISO 1)

The number of USV calls with peak frequencies <60 kHz (**Figure 6A**) showed a significant interaction between strain and postnatal day in both sexes. Across all strains, call frequencies were highest at PND 6 and decreased with age. In males, post hoc comparisons revealed that SD and SHHF pups emitted significantly fewer calls than Wistar pups on PND 6, whereas at PND 14, SHHF pups emitted more calls than SD pups (**Figure 6A1**). A similar pattern was observed in females, where SD pups emitted fewer calls than Wistar pups on PND 6 (**Figure 6A2**).

For call duration in the <60 kHz range (**Figure 6B**), significant effects of both strain and postnatal day were found in both males and females. SD pups consistently exhibited shorter call durations than SHHF and Wistar pups across postnatal days **(Figures 6B1 and 6B2**).

In the > 60 kHz frequency range (**Figure 6C**), call number increased with age across strains. In males, significant interactions and main effects of strain and postnatal day were found. On PND 14, SHHF pups emitted significantly more high-frequency calls than SD and Wistar pups (**Figure 6C1**). Female pups showed a similar developmental trend, with SHHF females emitting more high-frequency calls than the other strains at PND 14 (**Figure 6C2**).

Call duration > 60 kHz (**Figure 6D**) decreased over time with significant effects of strain. Wistar pups exhibited longer duration earlier in development (**Figure 6D1**). In females, a significant interaction was found, and SHHF and SD pups produced longer calls than Wistar on PND 14 (**Figure 6D2**).

##### Maternal potentiated USV calls (ISO 2)

For USV calls <60 kHz (**Figure 7A**), male pups showed a significant interaction between strain and postnatal day. Post hoc analyses revealed that SD and SHHF pups emitted fewer calls across postnatal days than Wistar pups. Additionally, Wistar and SHHF pups showed a progressive age-related decline in call number, whereas SD pups maintained relatively stable levels (**Figure 7A1**). A similar pattern was observed in females, with significant main effects and interaction, where SD and SHHF pups consistently emitted fewer calls than Wistar pups, and only Wistar and SHHF pups showed an age-related decrease (**Figure 7A2**).

Regarding call duration <60 kHz (**Figure 7B**), male pups showed a significant interaction and main effect of strain. Post hoc analysis revealed that at PND 9, SD pups exhibited shorter call duration than Wistar and SHHF pups (**Figure 7B1**). All strains showed an age-dependent increase in call duration. In females, significant effects of postnatal day and strain were found, with Wistar pups exhibiting slightly longer calls across the postnatal days (**Figure 7B2**).

For USV calls >60 kHz (**Figure 7C**), male pups showed a significant effect of postnatal day, strain, and their interaction. High-frequency calls increased with age, and post hoc test revealed that SD pups emitted more calls than Wistar pups at PND 9 and 14, with SHHF pups showing intermediate levels (**Figure 7C1**). In females, SHHF and Wistar pups emitted more high-frequency calls than SD pups across postnatal days (**Figure 7C2**).

Call duration>60 kHz (**Figure 7D**) in male pups showed significant main effects and interaction, with a general decline in duration with age. However, no specific strain differences reached significance in post hoc comparison (**Figure 7D1**). In female pups, no significant main or interaction effects were detected (**Figure 7D2**).

## 4. DISCUSSION

This study aimed to identify strain-dependent differences in early maternal behavior and pup development in Wistar, Sprague-Dawley (SD), and Spontaneously Hypertensive with Heart Failure (SHHF) rats. Maternal care was assessed during the first five postpartum days, revealing that SD dams showed lower frequencies of high-arched nursing and fewer behavioral transitions, despite increased licking behavior. SHHF dams spent more time in the nest without nursing and showed higher levels of self-grooming and nest-leaving during the dark phase. Network analyses revealed distinct organizational features: Wistar dams alternated consistently between nursing, nest presence, and brief absences; SD dams maintained a limited set of transitions concentrated in a few caregiving states; and SHHF dams showed narrower sequences with increasing OFF transitions across days. Pup development also varied by strain. SHHF pups had lower body weight, delayed eye opening, and slower reflex development, while SD pups showed earlier maturation in some reflexes but reduced vocalization duration. Ultrasonic vocalizations differed across isolation and maternal potentiation contexts, with Wistar pups emitting more calls early in life and SHHF pups showing a later increase in high-frequency calls.

### 4.1. Maternal behavior profile and strain differences

Maternal observations during the first five postpartum days revealed marked strain differences in the frequency, combination, and sequential organization of caregiving behaviors. Sprague-Dawley (SD) dams consistently displayed lower frequencies of high-arched nursing (HG) and less co-occurrence of licking while nursing (HG/L), despite exhibiting higher levels of isolated licking behavior, particularly during the light phase. This pattern suggests a more segmented caregiving style that may reflect less integrated responsiveness to pup needs. In contrast, SHHF dams showed high frequencies of OFF nest behavior, especially during the dark phase, suggesting possible dysregulation in balancing self-maintenance and pup care.

The transition network analysis provided further insight into the temporal and structural organization of maternal behavior. Wistar dams exhibited dynamic networks characterized by frequent transitions between high-arched nursing (HG), in-nest without nursing (DN), and OFF states (e.g., HG → DN → OFF → HG). This pattern suggests a diversified behavioral strategy that alternates consistently between active care, passive presence, and self-maintenance. In contrast, SD dams displayed more restricted networks, with fewer unique transitions and repeated cycling between DN and HG (e.g., DN → HG → DN), often without reaching OFF or incorporating pup-licking behaviors. Their networks were dominated by prolonged stays in passive states like DN, indicating reduced behavioral variability. SHHF dams showed more constrained and progressively narrowed networks, with increasing transitions to OFF states across days and fewer returns to active nursing.

From a developmental perspective, these patterns of behavioral transitions may represent distinct configurations of sensory scaffolding that shape the pup’s early experience. Different maternal sequences, such as alternations between nursing and licking or between presence and absence from the nest, may generate varied streams of tactile, thermal, and olfactory input. These sensory patterns unfold during a period of heightened neuroplasticity and may contribute to the differentiation and maturation of multiple neural circuits. As discussed in our previous work using the limited bedding and nesting paradigm (Pardo et al., 2024), disruptions in the temporal structure of maternal care may alter the ongoing maturation of experience-dependent systems guided by these sensory cues. In this context, the caregiving trajectories observed across strains may reflect distinct ways in which the maternal environment contributes to shaping neurodevelopmental outcomes in the offspring.

These strain-specific profiles likely reflect genetic and physiological differences that modulate maternal responsiveness. In both rats and mice, previous studies have shown that caregiving styles vary across strains and are linked to differences in emotional reactivity and behavioral outcomes in the offspring. In rats, Champagne et al. (2003) demonstrated that naturally occurring differences in licking/grooming among Long Evans dams are stably expressed across litters and associated with differences in stress reactivity and cognitive performance in the offspring, even though all mothers provided sufficient care to ensure survival and normal weaning weights. In mice, BALB/c and C57BL/6 strains differ in maternal licking frequencies and emotional reactivity, and these traits are associated with divergent offspring behavioral profiles even under cross-fostering conditions (Bagley et al., 2019; Priebe et al., 2005; Van Der Veen et al., 2008). In addition, Sprague-Dawley females are more prone than Wistar to develop maternal behavior in virgin sensitization paradigms involving repeated pup exposure (Jakubowski & Terkel, 1985). Beyond maternal care, Wistar and SD rats differ in their coping strategies and responsiveness to social and pharmacological challenges. For instance, Wistar rats show more active coping and reduced fear conditioning following social conflict (Walker et al., 2009), as well as greater behavioral reactivity to opioid modulation during adolescent social play (Manduca et al., 2014).

Susceptibility to early-life challenges also varies across strains. Our group previously reported that the limited bedding and nesting (LBN) paradigm induces more pronounced changes in maternal care and offspring development in BALB/c mice than in C57BL/6 (Pardo et al., 2023). Likewise, Neeley et al. (2011) showed that prenatal stress alters hippocampal gene expression in a strain-dependent manner, with Sprague-Dawley and Lewis rats exhibiting more pronounced changes in Bdnf, Grin2b, and Tnfα expression compared to Fischer rats, which showed a more stable molecular profile. In terms of neurodevelopmental vulnerability, Zmarowski et al. (2012) found that SD rats are more susceptible than Wistar Han to prenatal neurotoxic exposure, showing greater cognitive impairments and hippocampal abnormalities. These findings emphasize that strain differences extend beyond behavioral tendencies to include neurobiological and developmental responses to environmental perturbations.

Particularly novel in this study is the characterization of maternal behavior in the SHHF strain, which had not been previously described in this context. The hypertensive and heart failure-prone background of SHHF dams may contribute to their distinct maternal strategy, paralleling findings in Wistar-Kyoto (WKY) dams. Alves et al. (2022) reported that WKY dams, considered a model of depressive-like vulnerability, increased contact with their pups after maternal separation but showed reduced affiliative behaviors such as licking and grooming. These caregiving differences were associated with impaired cognitive outcomes in the offspring. Similarly, studies in SD rats have shown that naturally occurring variation in maternal licking is associated with long-term differences in offspring emotional behavior and monoamine signaling in the prefrontal cortex (Masís-Calvo et al., 2013).

Together, these findings support the notion that strain-dependent maternal profiles are not merely behavioral variants but reflect biologically grounded differences in caregiving organization, emotional responsiveness, and offspring neurodevelopment. By systematically characterizing maternal behavior in SHHF dams and comparing it with that of two widely used laboratory strains, this study provides novel insights into the diversity and functional relevance of early caregiving strategies in rodents.

### 4.2. Developmental physical milestones and reflexes profile of pups

Strain differences in pup development extended beyond maternal care, encompassing both somatic growth and the maturation of early reflexes. SHHF pups of both sexes exhibited reduced weight gain from early postnatal days (PND 6 in females, PND 10 in males), as well as delayed eye opening compared to Wistar and SD pups. In contrast, SD pups reached this milestone earlier, aligning with previous descriptions of their early postnatal locomotor development (Altman & Sudarshan, 1975). The SHHF profile suggests a slower pace of somatic maturation, which may reflect underlying physiological vulnerabilities associated with their hypertensive and cardiometabolic background.

The emergence and efficiency of neurodevelopmental reflexes further differentiated the strains. SHHF pups displayed longer latencies in completing the surface righting and negative geotaxis responses, indicating reduced postural control and sensorimotor coordination. Additionally, auditory startle responses were notably weaker in SHHF pups at PND 14, suggesting a delayed maturation of auditory pathways or arousal mechanisms. Although all strains improved with age, SHHF pups consistently lagged behind. In the grasp reflex, there were no significant differences among males; however, SHHF and Wistar females outperformed SD females at PND 6.

These findings are consistent with evidence that early sensorimotor reflexes are sensitive indicators of neurological maturation in rodents (Fox, 1965; Heyser, 2003; Nguyen et al., 2017). Reflexes such as righting, geotaxis, and startle rely on the functional integrity of brainstem and spinal circuits, and delays in their expression may signal slower development of sensorimotor integration or incomplete myelination. Supporting this, Smirnov and Sitnikova (2019) demonstrated that early sensory deprivation, through transient whisker removal, delays the emergence of multiple developmental milestones in rat pups, including eye opening, locomotor coordination, grooming, and rearing. Their findings highlight the role of early sensory input in organizing sensorimotor development and suggest that even minor perturbations during critical periods can shift the timing of functional emergence. In this context, the reflex delays observed in SHHF pups may reflect not gross dysfunction but a slower progression of sensory integration systems, potentially shaped by strain-specific physiological characteristics.

Importantly, these differences in SHHF were not associated with the absence of responses or gross deficits, but rather slower and less efficient performance. Whether this reflects a transient maturational delay or a more persistent strain-specific neurodevelopmental phenotype remains to be determined. Previous studies have shown that early adversity can modulate the timing of physical and reflex milestones depending on genetic background. For example, in our previous work, we demonstrated that limited bedding and nesting (LBN) exposure delayed eye opening and impaired weight gain in BALB/c and C57BL/6 mouse pups, with effects differing by strain and sex (Pardo et al., 2023). Such evidence supports the idea that early developmental indicators can reveal subtle neurobiological vulnerabilities.

Although our study did not track long-term outcomes, the profile observed in SHHF pups suggests a distinct maturational trajectory. Longitudinal studies are needed to determine whether these early sensorimotor patterns can predict later behavioral or cognitive changes, as suggested by developmental models that link early delays with deficits observed during adolescence (Abramova et al., 2021; Baharnoori et al., 2012). Such studies will be crucial in determining whether these early profiles are indicative of long-term behavioral or physiological vulnerabilities.

### 4.3. Ultrasonic vocalization profile of pups

The ultrasonic vocalizations (USVs) emitted by pups during isolation provide a valuable window into the early development of affective communication and caregiver–infant dynamics. Across all three strains, the majority of USVs during early postnatal days occurred at frequencies below 60 kHz, which are commonly associated with states of distress, low thermal regulation, and the need for maternal contact (Boulanger-Bertolus et al., 2017; Hashimoto et al., 2007). As pups aged, there was a gradual increase in the proportion of high-frequency calls (>60 kHz), particularly around PND 9–14, although the timing and intensity of this shift differed between strains. These high-frequency calls, typically shorter in duration, are thought to reflect increased motor activity or exploratory drive and may function as developmentally emergent signals within the affective or social communication system (Stark et al., 2020). In our data, Wistar pups displayed the expected developmental pattern, with high rates of <60 kHz calls in early stages and a subsequent emergence of >60 kHz calls. SHHF pups, by contrast, showed a delayed but marked increase in >60 kHz calls by PND 14, while SD pups maintained a flatter profile across frequencies and ages. These observations suggest both a conserved ontogenetic transition and strain-specific differences in the maturation of infant communication patterns.

These findings align with the well-established phenomenon of maternal potentiation, whereby brief contact with the dam enhances subsequent vocal output during a second isolation (Hofer et al., 1998; Shair, 2014). This response is considered a behavioral marker of filial bonding, dependent on sensory cues and neurochemical signaling pathways that underlie the social memory of the dam (Rohitsingh et al., 2011; Shair, 2014). The reduced potentiation observed in SD and SHHF pups may reflect either a delay or disruption in the development of this affiliative signaling system.

Importantly, USVs are not merely passive responses to stress but form part of a bidirectional regulatory system between mother and infant. These vocalizations induce maternal retrieval, licking, and nest-building behaviors (Hashimoto et al., 2001; Wöhr & Schwarting, 2008) and are modulated by prior maternal care (Brunelli et al., 2015; Zimmerberg et al., 2003). Moreover, variation in USV profiles among individuals and strains has been linked to long-term differences in emotional reactivity and social behavior (Brunelli & Hofer, 2007; Granata et al., 2022). From a developmental perspective, the ontogeny of USVs reflects both physiological maturation and environmental modulation. Typically, USVs peak between PND 6–9 and decline by PND 14–21 (Hashimoto et al., 2007; Stark et al., 2020). The delayed increase in vocalizations observed in SHHF pups suggests a possible shift or prolongation in this developmental trajectory. This pattern could indicate either neurodevelopmental immaturity, as proposed in models of prenatal disruption (Gulia et al., 2014), or altered sensory processing, potentially related to the SHHF strain’s cardiovascular and autonomic phenotype.

While traditionally interpreted as distress signals, alternative hypotheses propose that USVs may arise from physiological mechanisms such as the abdominal compression reaction during respiratory effort or thermoregulation (Blumberg et al., 2000; Blumberg & Alberts, 1991). These interpretations suggest that variation in USVs might not always reflect emotional distress *per se*, but rather differences in internal state regulation. This perspective is particularly useful for understanding cases like SHHF, where vocal output increases despite lower maternal potentiation.

Finally, the acoustic structure of USVs also carries functional significance. Boulanger-Bertolus et al. (2017) distinguished between lower-frequency “distress” calls and higher-frequency calls potentially linked to movement or exploratory states. The predominance of high-frequency calls in SHHF pups may reflect altered sensory-affective integration, further supported by evidence that USVs predict adolescent social behavior and are linked to neuropeptide expression (Granata et al., 2022).

Taken together, our findings support the notion that strain-dependent differences in USV profiles during isolation and maternal potentiation paradigms reflect distinct developmental trajectories of the affective communication system. These differences likely result from the interplay between genetic background, physiological maturation, and early maternal care, underscoring the value of USVs as a sensitive, multi-dimensional tool for studying early socio-affective development in rodent models and their relevance for human neurodevelopmental disorders.

### 4.4. Methodological considerations and translational relevance

When interpreting the current findings, it is important to consider methodological factors. All animals were evaluated at a high-altitude location (approximately 3300 meters above sea level), where mild chronic hypobaric hypoxia may affect physiological development. Although all strains were equally exposed to this environment, strain-specific differences in oxygen sensitivity, metabolic rate, or cardiovascular function could have influenced their developmental trajectories. For instance, altitude-related hypoxia may interact with the underlying hypertensive and cardiopulmonary phenotype of SHHF rats, potentially impacting growth rate, reflex development, and vocal communication. Future studies that compare rearing conditions in highland and lowland environments would help clarify the contributions of genetics and environment to these outcomes.

This study was conducted at an elevation of 3300 meters above sea level, in a hypobaric environment that introduces a chronic hypoxic challenge during critical periods of maternal and early postnatal development. Evidence shows that high-altitude exposure affects rodent physiology and behavior in species- and strain-specific ways. For instance, rats and mice differ in their strategies of acclimatizing to hypoxia, with mice displaying greater ventilatory and mitochondrial plasticity and rats relying more on hematological compensation (Arias-Reyes et al., 2021). Additionally, research indicates that maternal behavior is sensitive to altitude-induced hypoxia. Native deer mice exposed to chronic hypoxia during lactation show reduced licking behavior and altered maternal motivation (Robertson & McClelland, 2021). Furthermore, long-term colonies of rats and mice bred at high altitude display distinct ventilatory, pulmonary, and metabolic phenotypes compared to those at sea level (Jochmans-Lemoine et al., 2015). These findings emphasize the importance of considering altitude as a modulatory variable in behavioral and developmental studies using rodents.

This study emphasizes the importance of examining multiple strains in developmental neuroscience from a comparative perspective. Even when raised under the same conditions, Wistar, Sprague-Dawley, and SHHF rats showed significant differences in maternal behavior, growth, the emergence of reflexes, and patterns of ultrasonic vocalization. These findings highlight that baseline neurodevelopmental processes can vary across different strains, which may greatly affect experimental outcomes. Notably, identifying strains with unusual or extreme characteristics, such as the unique maternal behavior and altered vocalization patterns observed in SHHF rats, provides valuable opportunities to study models of vulnerability that occur naturally, without the need for external interventions.

Finally, this work provides novel descriptive data on the SHHF strain in the context of early neurobehavioral development, extending its translational utility beyond cardiovascular and metabolic research. Given the growing interest in the developmental origins of health and disease, our findings suggest that SHHF rats may represent a promising model for investigating the early-life interplay between genetic predisposition, caregiving quality, and the maturation of affective and physiological systems. More broadly, these results support the integration of strain selection, environmental context, and behavioral phenotyping as core elements in preclinical models of neurodevelopment.

### 4.5. Conclusion

This study presents a comprehensive comparative analysis of early maternal behavior and neurodevelopmental milestones in three commonly used rat strains: Wistar, Sprague Dawley (SD), and Spontaneously Hypertensive Heart Failure (SHHF). By examining behavioral transitions, reflex development, somatic growth, and ultrasonic vocalizations, we identified consistent strain-dependent profiles in maternal strategies and the developmental trajectories of their offspring. These differences were observed under standardized laboratory conditions, reinforcing the significant role of genetic background as a fundamental factor influencing early caregiving and neurobehavioral development. Our findings emphasize the importance of integrating both conventional and qualitative analyses of behavioral measures, such as maternal behavior transition networks and ultrasonic vocalization patterns, to capture the complexities of early-life caregiving and infant communication. The characterization of SHHF rats, in particular, provides new insights into how a genetic predisposition to cardiovascular and metabolic dysfunction may impact maternal behavior and the early development of emotional systems. Overall, this research underscores that strain selection is not a secondary methodological consideration but a fundamental variable in neurodevelopmental research. Future studies leveraging this variability may deepen our understanding of early vulnerability and resilience, offering new avenues for translational models of health and disease.

## Author contribution

**Grace E. Pardo:** conceptualization, methodology, validation, formal analysis, investigation, resources, data curation, visualization, writing–original draft, writing– review and editing, supervision, project administration, funding acquisition. **Alondra Casas Pary**: visualization, investigation, validation, data curation. **Maria Esther Gutierrez Ccori**: investigation, data curation. **Mabel Choque Aguilar**: investigation, data curation. **Lucero B. Cuevas**: visualization, software. **Enver M. Oruro**: validation, visualization, software, writing–review and editing. **Luis F. Pacheco-Otalora**: conceptualization, resources, writing–review and editing.

